# Zero-shot ecological annotation of microbial genomes with myLLannotator accelerates scientific discovery

**DOI:** 10.64898/2026.01.18.700140

**Authors:** Alyssa Lu Lee, Arya Sharma, Rohan Maddamsetti

**Affiliations:** Institute for Quantitative Biomedicine, Rutgers University, New Brunswick, NJ; Graduate Program in Molecular Biosciences, Rutgers University, New Brunswick, NJ; Department of Biochemistry and Microbiology, Rutgers University, New Brunswick, NJ

## Abstract

Large language models (LLMs) are promising scientific assistants, but many models remain inaccessible for resource-constrained scientists. To address this, we present myLLannotator, a python package for metadata annotation built on llama3.2-3B. myLLannotator labels 18,000+ genomes in ~2 hours on a laptop, reproducing a discovery involving substantial manual annotation. We use myLLannotator to discover that duplicated genes are depleted in endosymbiotic bacteria, demonstrating its power to accelerate hypothesis testing and discovery.

## MAIN

Public biological databases contain a wealth of hidden insights, but researchers seeking to explore data and test hypotheses are often hindered by the substantial time and effort required to manually annotate samples based on inconsistent or missing metadata. Furthermore, even when data are annotated with standardized semantic terms from a well-designed ontology^1^, researchers still need to map the terms of a given ontology onto their particular scientific question, which may require combining or splitting keywords into relevant categories.

In principle, large language models (LLMs) can flexibly address this problem, as they excel at completing natural language tasks based on unstructured textual prompts^2,3^. Indeed, benchmarking shows that LLMs can accurately annotate cell types and gene sets based on user prompts^4-6^. Despite exciting ongoing developments, it is unclear whether LLM assistants will accelerate discovery for most scientists, for two reasons. First, most of these tools are either inaccessible to the broader scientific community^7-10^, are based on closed-source LLMs that require a commercial subscription^4^, or require substantial computational resources for inference. This lack of access poses a significant barrier to many, if not most, scientists and students across the globe. Second, the cost of conducting experiments to test hypotheses far outweighs the effort required for hypothesis generation, even if assisted by an LLM^8,11^. Therefore, what is needed is an experiment that would test whether an average human researcher with limited resources can use an LLM to make new discoveries that would be otherwise be cost-prohibitive in terms of time and effort.

To answer this question, we present myLLannotator, a python package built on the lightweight open-source LLM llama3.2-3B^12^, and use it to classify tens of thousands of microbial isolates into researcher-defined categories based on unstructured free-text host source, isolation source, and species metadata fields. As a positive control, we show that zero-shot ecological annotation with myLLannotator accurately reproduces a key result from previously published work. We then show that myLLannotator allows researchers to quickly test new hypotheses by rapidly reannotating thousands of genomes in about 2 hours, using basic computational resources such as a laptop computer.

In previous work, we classified 18,938 bacterial genomes from NCBI RefSeq into 7 ecological categories to test the hypothesis that duplicated antibiotic resistance genes would be enriched in environments associated with antibiotic use^13^. The original analysis required substantial manual annotation taking days of effort, which was then codified in a python script to categorize the ecological provenance of microbial genomes based on a very large set of handcrafted pattern-matching rules against the isolation source and host source metadata found in RefSeq genome annotation files^13^. Although there are large databases of ecological annotation for microbial genomes, such as the proGenomes database^14^ and the Integrated Microbial Genomes & Microbiomes (IMG/M) database^15^, we found such existing resources insufficient for our purposes. The ecological annotations in the proGenomes database are hidden behind a frustrating web search interface— the full underlying annotation metadata tables are inaccessible for most users— and downloadable annotations are only available for a select number of representative genomes. By contrast, the IMG/M database has a well-designed interface for downloading ecological metadata, but users are constrained to use the pre-existing labels and categories in the database, which may not necessarily be relevant for the hypotheses at hand. Furthermore, combining metadata from one database (such as IMG/M) with data from a different database (genomes from NCBI RefSeq) raises the basic challenge of getting disparate datasets into a consistent format for merging sets of genome IDs and annotations—a common and time-consuming task requiring careful attention and manual effort. We hypothesized that we could replace our handcrafted ecological annotation rules with a short and simple LLM prompt describing the rationale behind our original annotations (Methods).

We use an alluvial diagram^16,17^ to visualize how our original annotations^13,18^ map onto the annotations produced by myLLannotator (Figure 1A). Including unannotated genomes (those with insufficient metadata for annotation), 15,062 out of 18,938 genome annotations matched, achieving 79.5% accuracy. Excluding unannotated genomes, 15,062 out of 18,062 genome annotations matched, achieving 83.4% accuracy.

**Figure 1:**
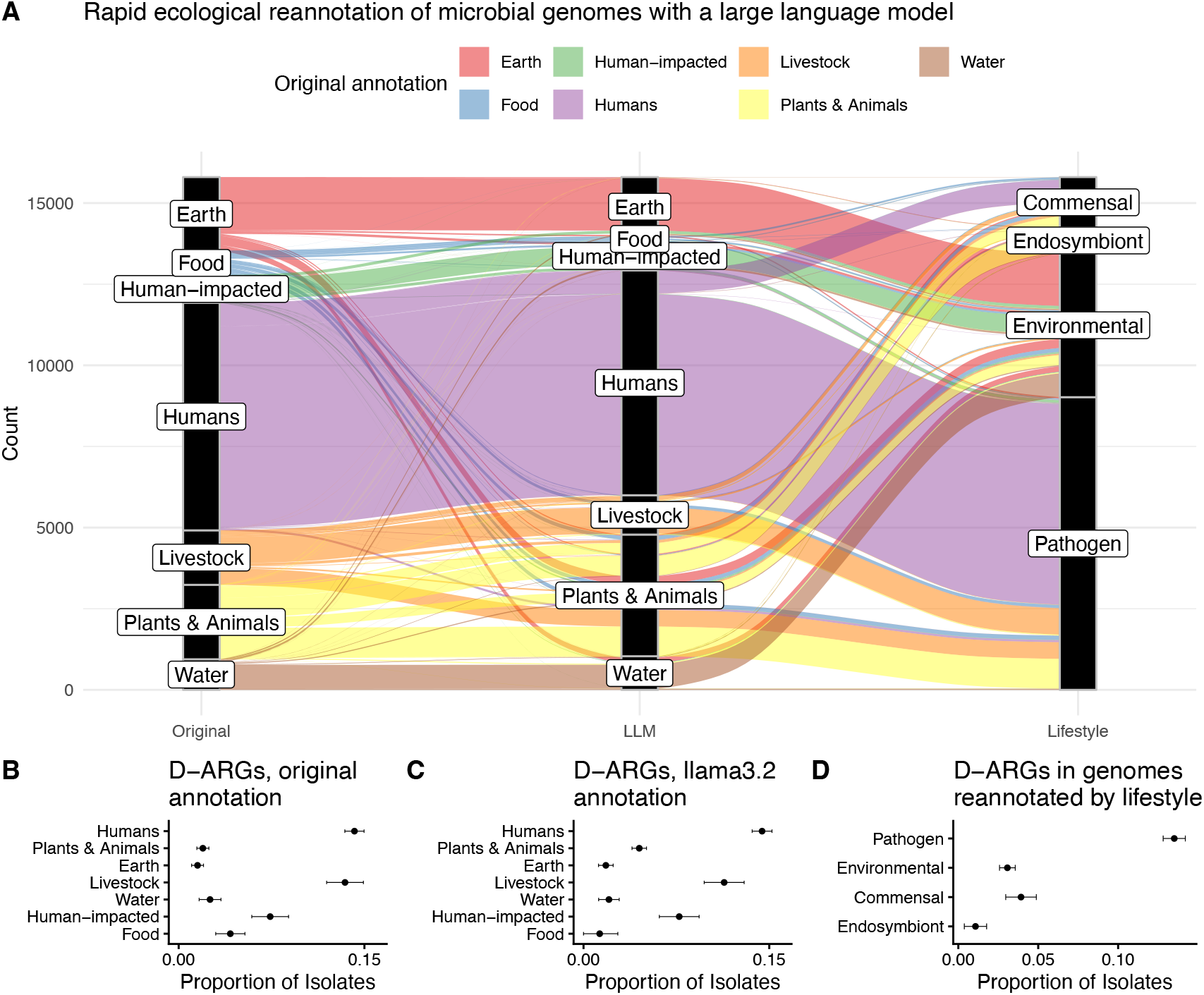
Zero-shot classification of microbial genomes with myLLannotator. A) An alluvial diagram shows that zero-shot ecological annotations produced by llama3.2 are highly consistent with semi-manual annotations from Maddamsetti et al. (2024) and are easily remapped into pathogen, endosymbiont, commensal, and environmental categories. B) Key result reported by Maddamsetti et al. 2024, based on semi-manual annotation: duplicated antibiotic resistance genes (D-ARGs) are enriched in microbial isolates from humans and livestock. C) Fully automated LLM annotation with llama3.2 reproduces the key result reported by Maddamsetti et al. 2024: duplicated antibiotic resistance genes (D-ARGs) are enriched in microbial isolates from humans and livestock. D) Zero-shot reannotation of bacterial isolates with llama3.2 reveals that D-ARGs are enriched in pathogens compared to commensal, endosymbiont, and environmental isolates.

Our original result, showing that duplicated antibiotic resistance genes (D-ARGs) are enriched in bacteria isolated from humans and livestock, is shown in Figure 1B. The same analysis, conducted on samples annotated by llama3.2, is shown in Figure 1C. A comparison of panels B and C shows that llama3.2 accurately reproduces the key result reported by Maddamsetti et al. (2024).

Automated annotation also enabled us to rapidly test a new hypothesis. First, we used myLLannotator to classify samples by host lifestyle (pathogen, commensal, environmental, or endosymbiont) based on isolation source, host, and species name metadata (Figure 1A). Reannotation was practically effortless, taking a simple prompt (Methods) and 2 hours and 15 minutes of runtime on a 2021 M1 Macbook Pro laptop. Without the LLM, complete reannotation of this dataset would have required substantial time and effort. The reannotation reveals that D-ARGs are enriched in pathogenic isolates compared to environmental, commensal, and endosymbiont isolates (Figure 1D). Combined with the observation that D-ARGs are enriched in humans and livestock (Figure 1B), this finding supports the claim made by Maddamsetti et al. 2024 that higher rates of antibiotic exposure drive the evolution of duplicated antibiotic resistance genes. The goal of this reannotation, however, was to test a different idea: from our previous work, we had noticed that bacteria isolated from plants and animals had fewer duplicated genes (D-genes) compared to bacteria in the other categories (Figure 2A) and had fewer single-copy antibiotic resistance genes (S-ARGs) (Figure 2B). From our manual annotation, we knew that many endosymbiotic bacteria isolated from insects were classified in this category. Since is known that many endosymbiotic bacteria have highly reduced genomes^19,20^, we hypothesized that the signals of fewer D-genes and fewer S-ARGs in plant- and animal-associated bacteria in Figure 2A and Figure 2B might be driven by endosymbiotic bacteria. Indeed, we find strong evidence for this hypothesis: our reannotation shows that endosymbiotic bacteria are depleted in D-genes (Figure 2C) as well as S-ARGs (Figure 2D) by comparison with bacteria annotated as pathogens, environmental isolates, and commensals.

**Figure 2:**
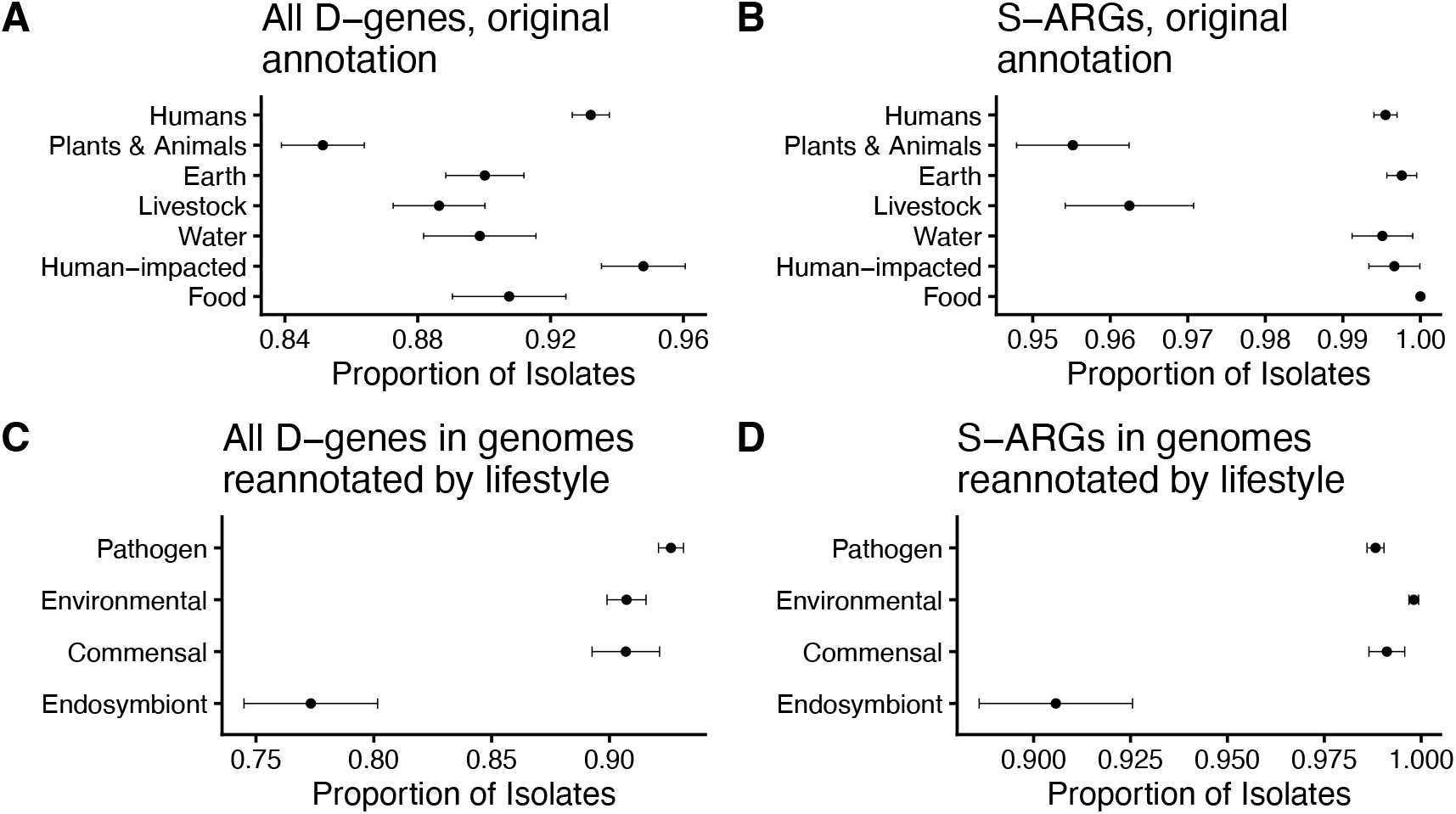
Zero-shot ecological reannotation of microbial genomes with myLLannotator accelerates scientific discovery. A) Analysis of duplicated genes (D-genes) from Maddamsetti et al. 2024 shows a weak signal of fewer D-genes in plant- and animal-associated bacteria. B) Analysis of single-copy antibiotic resistance genes (S-ARGs) from Maddamsetti et al. 2024 shows a signal of fewer S-ARGs in bacteria isolated from livestock and plants and animals. C) Zero-shot reannotation of microbial genomes with llama3.2 shows that duplicated genes (D-genes) are depleted in endosymbiotic bacteria. D) Zero-shot reannotation of microbial genomes with llama3.2 shows that single-copy antibiotic resistance genes (S-ARGs) are depleted in endosymbiotic bacteria.

Most existing studies on LLMs for data annotation focus on accuracy against artificial benchmarks^21^. By contrast, our paper focuses on the benchmark that scientists primarily care about: whether a tool can accelerate their ability to make discoveries. Our results conclusively demonstrate that current open-source LLMs such as llama3.2 dramatically accelerate the ability of researchers to generate biological insights, by removing the need for extensive manual annotation before testing a promising hypothesis on large-scale public datasets.

Our python package, myLLannotator, is simple and fast, taking two hours to annotate ~18,000 samples, and is based on a tiny open-source LLM that has only 3 billion parameters and runs quickly on modern laptop computers. We emphasize that myLLannotator can be applied to any scientific dataset lacking standardized metadata, from microbiome samples to plant, animal, and medical specimens. In addition, myLLannotator is compatible with open-source LLMs beyond llama3.2, some of which may have greater capabilities. We encourage researchers to adapt our code for their own specific use cases. We anticipate that the method followed here, of using an LLM to quickly annotate samples to explore patterns in data and rapidly test hypotheses, will be broadly adopted across the biological sciences.

## Methods

### Input data

The input data files for this analysis are available at: https://rutgers.box.com/v/myLLannotator-data.

### Ecological annotation

A python 3.12 script called *annotator*.*py* was used to reannotate the genomes analyzed by Maddamsetti et al. (2024), using the llama3.2-3B LLM, installed and called through the Ollama python API (https://docs.ollama.com/).

We used the following prompt to reannotate genomes based on the annotation rules used by Maddamsetti et al. (2024):

> “You are an annotation tool for labeling the environment category that a microbial sample came from, given the host and isolation source metadata reported for this genome. Label the sample as one of the following categories: “+ categories_str +” by following the following criteria. Samples from a human body should be labeled ‘Humans’. Samples from domesticated or farm animals should be labeled ‘Livestock’. Samples from food should be labeled ‘Food’. Samples from freshwater should be labeled ‘Freshwater’. Samples from a human-impacted environment or an anthropogenic environmental source should be labeled ‘Anthropogenic’. Samples from the ocean, including the deep ocean should be labeled ‘Marine’. Samples from anoxic sediments, including aquatic sediments should be labeled ‘Sediment’. Samples from domesticated plants and crops should be labeled ‘Agriculture’. Samples from soil, including farm soil should be labeled ‘Soil’. Samples from extreme terrestrial environments (extreme temperature, pH, or salinity) should be labeled ‘Terrestrial’. Samples from non-domesticated or wild plants should be labeled ‘Plants’. Samples from non-domesticated or wild animals, including invertebrates, and also including protists and fungi even though these are not strictly animals should be labeled ‘Animals’. Samples that lack enough information in the host metadata and isolation source metadata provided or have missing or incomplete information in these fields should be labeled ‘NA’. Give a strictly one-word response that exactly matches of these categories, omitting punctuation marks.”

We used the following prompt to reannotate genomes as Pathogen, Commensal, Endosymbiont, or Environmental:

> “You are an annotation tool for labeling microbial samples, given its species names and the host and isolation source metadata reported for this sample. Label the sample as one of the following categories: “+ categories_str +” by following the following criteria. Samples from blood, organs, or diseased sites or disease should be labeled ‘Pathogen’. Samples from normal or healthy plant or animal or human body sites should be labeled ‘Commensal’. Samples from the environment should be labeled ‘Environmental’. Samples that are obligate endosymbionts, based on its species and isolation from its obligate plant or animal host and isolation source should be labeled ‘Endosymbiont’. Samples that lack enough information in the host metadata and isolation source metadata provided or have missing or incomplete information in these fields should be labeled ‘NA’. Give a strictly one-word response that exactly matches these categories, omitting punctuation marks.”

### Data analysis and figure generation

Data analysis was conducted with an R 4.2 script called simple-ARG-duplication-analysis.R, which generated all figures. Run times were measured on a 2021 Macbook Pro with an M1 Max CPU and 64GB of RAM.

## Data Availability

Accessions for the 18,938 complete bacterial genomes from NCBI RefSeq analyzed in this work are listed in Supplementary Data 3 of Maddamsetti et al. (2024) (reference 13). A minimum dataset necessary to interpret, verify and extend the research in this article is available at: https://rutgers.box.com/v/myLLannotator-data.

## Code Availability

A Github repository containing all code sufficient to reproduce this work, including statistics and figures, is available at https://github.com/alyssa-lee/myLLannotator. myLLannotator is also available at: https://pypi.org/project/myllannotator/

## Acknowledgements

This work was partially funded by the New Jersey Agricultural Experiment Station (startup funds to R.M.) and by the Rutgers Health BMIHAI Pilot Grant Program.

